# Evolution of the Jawed Vertebrate (Gnathostomata) Stomach through Gene Repertoire Loss: Findings from Agastric Species

**DOI:** 10.1101/2024.12.02.626498

**Authors:** Jackson Dann, Frank Grützner

**Affiliations:** School of Biological Sciences, The University of Adelaide, SA, 5005, Australia; Robinson Research Institute, The University of Adelaide, SA, 5005, Australia

## Abstract

Independent and convergent gene loss, such as the loss of gastric enzymes (*Pga*, *Pgc*) and proton pump subunits (*Atp4a*, *Atp4b*) in monotremes and polyphyletic teleost species have been associated with the agastric phenotype – stomachs lacking gross morphology, gastric glands and acid secretion. The loss of the key gastric hormone gastrin (*Gast*) has been observed in monotremes, but not corroborated in agastric teleosts. Furthermore, it is unclear how this loss affects the evolution and selection of the native receptor (*Cckbr*), gastrin releasing peptide (*Grp*) and receptor (*Grpr*), or if there are further gene losses associated with the agastric phenotype. We investigated the evolution, selection and conservation of the gastrin physiological pathway and novel gastric genes in a broad range of gastric and agastric vertebrates. Our analyses showed no correlation between the losses of *Gast* and *Grpr* with the agastric phenotype in vertebrate lineages. We identified genes implicated in gastric mucosal maintenance and development (*Gkn1*, *Gkn2*, *Tff1*, *Tff2*, *Vsig1*, *Anxa10*) that are lost in agastric species except the echidna, which retained some of them (*Gkn1*, *Tff2*, *Vsig1*). Our findings demonstrate a convergent repertoire of gastric genes lost in agastric species, partial retention of gastric functionality in the echidna, and the diversified roles of gastrin pathway constituents in vertebrate lineages.

## Introduction

The stomach is a highly conserved and integral organ throughout jawed vertebrate (Gnathostomata) evolution as evidenced by conserved developmental pathways dating back to protochordate species (Nakazawa et al., 2013; Nakayama et al., 2017). Though morphological features can vary widely between taxa, comparable components include storage compartments (enlargement of the lumen) and acid-secreting glandular epithelium (Koelz, 1992; Karasov, Douglas & Ronald, 2013). Despite gastric features being nearly ubiquitous, a unique gastric phenotype has evolved over 15 times independently in ray-finned fish (Actinopterygii), and at least twice in lobe-finned fish (Sarcopterygii): once in lungfish and once in monotremes. This gastric phenotype – hereby known as agastric – is typified by an absence of acidic luminal contents, loss of glandular epithelium and a significant reduction in overall size and gross morphology (Wilson & Castro, 2010; Castro et al., 2014).

Not only is the agastric phenotype convergent morphologically between species, but agastric species share the concomitant loss of genes for gastric enzymes (*Pga*, *Pgc*) and proton pump subunits (*Atp4a*, *Atp4b*). In addition to this, monotremes have had an inactivation of the NK3 homeobox 2 (*Nkx3.2*) and have lost gastrin (*Gast*): a key gastric hormone regulated by the upstream neuropeptide gastrin-releasing peptide (*Grp)* and its native receptor (*Grpr*), which when released and binds the native cholecystokinin B receptor (*Cckbr*) potentiates gastric acid secretion (Ordoñez et al., 2008; Schubert & Peura, 2008; Castro et al., 2014; Zhou et al., 2021; Dann et al., 2024). Given the heterogeneity of the gastric epithelium in cell type (e.g. parietal cell, mucous cells, stem cells) and function (e.g. acidic digestion, immunity, hormone release), and various physiological and intracellular signalling pathways regulating gastric development and function, many more genes are expected to be affected by this drastic shift in gastric anatomy and physiology (Kim & Shivdasani, 2016; Agace & McCoy, 2017).

Recent discoveries in vertebrate developmental genetics and genomics have expanded the repertoire of genes implicated in gastric development and function. These have included the constituents of the gastrokine and trefoil factor families (*Gkn1*, *Gkn2*, *Gkn3*, *Tff1*, *Tff2*); secreted products of gastric mucosal epithelia with both independent and synergistic anti-inflammatory, tumour-suppressing and anti-apoptotic roles (Menheniott, Kurklu & Giraud, 2013; Hoffman, 2020). As well, *Vsig1* – a junction adhesion molecule expressed highly in gastric epithelia – and annexin A10 (*Anxa10*) – a component of the phospholipid-binding protein annexin superfamily – have displayed restricted gastric epithelial expression and been implicated in squamous versus glandular epithelial differentiation (Lu et al., 2011; Oidovsambuu et al., 2011). Despite some extensive work on expression patterns (particularly in gastrokine and trefoil factor families), and a recent work by Kato et al., (2024) – which noted the absence of *Vsig1* in agastric teleost lineages and the platypus – it is still currently unclear what emergent properties (i.e. gene ontologies) arise from the list of lost genes, and whether these additional genes are conserved in agastric vertebrate lineages (Jiang, Lossie & Applegate, 2011; Geahlen et al., 2013).

To better understand the evolution of the agastric phenotype in the Actinopterygii and Sarcopterygii clades, we explored the evolution, selection and loss of the gastrin physiological pathway along with novel gastric gene sets in the genomes of gastric and agastric taxa.

## Materials and Methods

### Sequence retrieval and synteny analysis

Sequences were obtained from evolutionary representative taxa of tetrapods (placental mammals, marsupials, monotremes, reptiles), Sarcopterygii (lobe-finned fish) and Actinopterygii (ray-finned fish). Annotated protein sequences were downloaded from NCBI GenBank (https://www.ncbi.nlm.nih.gov/genbank/) and, where no annotations were present, searched for through the BLAST tool using homologous genes from closely-related species (https://blast.ncbi.nlm.nih.gov/Blast.cgi). Synteny analysis was performed in the NCBI genome data viewer tool (https://www.ncbi.nlm.nih.gov/genome/gdv/). See Supplementary file 1 for complete sequence and genome information.

### Sequence alignments and phylogenies

Multiple sequence alignments were performed using the Clustal Omega plugin in Geneious 11.0.14+1 except for deleted / pseudogenised monotreme genes (Geneious, 2022; Sievers et al., 2011). Protein maximum likelihood phylogenies were then constructed using the IQ-TREE web server (http://iqtree.cibiv.univie.ac.at/) tree inference tool with standard settings and 1000 bootstrap replicates (Trifinopoulos et al., 2016). Substitution models derived from Bayesian information criterion and alignment outgroups can be found in Table S1.

### Selection analysis (Ka/Ks ratios)

To detect signatures of selection, amino acid alignments and corresponding open reading frame DNA alignments were converted into PAML format codon alignments using PAL2NAL v 14 (Mikita, Torrents & Bork, 2006). Codon alignments along with their corresponding phylogenies in Newick format were then input into EasyCodeML v1.41 using the preset branch model with multiples models of foreground branch selection for each lineage and likelihood ratio tests (LRTs) to assess model significance (Gao et al., 2019). EasyCodeML outputs can be found in Table S2.

## Results

### Conservation and loss of the gastrin release pathway and novel gastric gene repertoire in gastric and agastric jawed vertebrates (Gnathostomata)

Synteny analysis of gastrin (*Gast*), the native gastrin receptor (*Cckbr*), the gastrin-releasing peptide (*Grp*) and its receptor (*Grpr*) shows conserved syntenic blocks in ray- and lobe-finned fish species with recurring differences of flanking loci between lineages (Figure 1A). Gastrin was lost in two of the three agastric species (pufferfish and west African lungfish), but conserved in the agastric zebrafish. Additionally, the gastrin-releasing peptide receptor (*Grpr*) was absent from conserved genomic neighbourhoods in the gastric three-spined stickleback (*Gasterosteus aculeatus*). The highly-conserved syntenic blocks of tetrapod species corroborated the previously-noted gene losses of gastrin in both monotreme species (Ordoñez et al., 2016), with our analyses outlining an unexpected absence of the *Grpr* locus in the platypus (Figure 1B). Interestingly, these findings suggest that the loss of gastrin does not correlate with the presence of the agastric phenotype, and, that the loss of gastrin does not correlate with the loss of *Grpr*.

**Figure 1:**
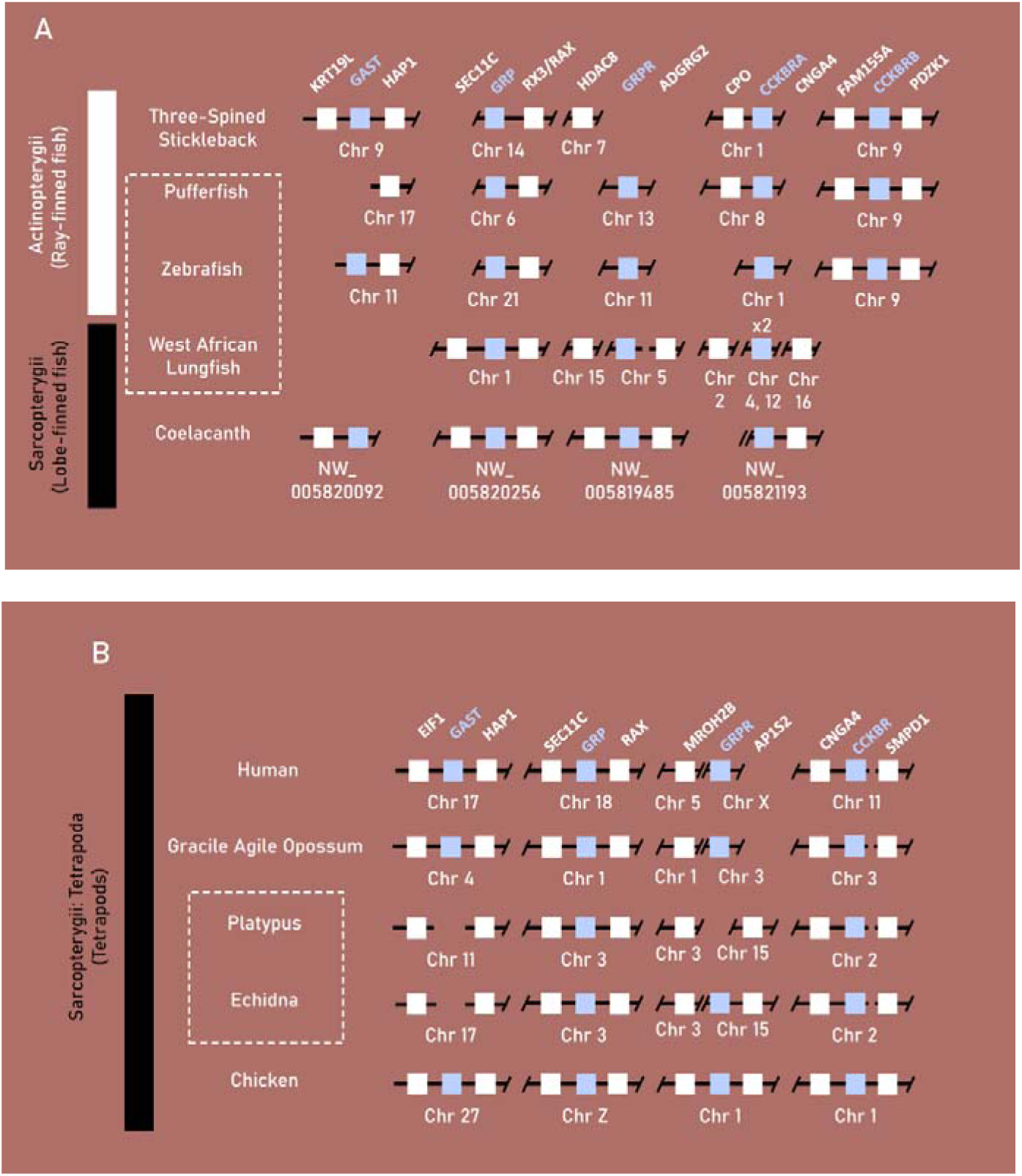
Synteny plots of gastrin (*Gast*), gastrin-releasing peptide (*Grp*), gastrin-releasing peptide receptor (*Grpr*) and cholecystokinin receptor B (*Cckbr*) in select vertebrate species. **A**, synteny plot of gastric genes lost or pseudogenised in the agastric species of the zebrafish (*D. rerio*) and West-African Lungfish (*P. annectens*) compared to the gastric three-spined stickleback (*G. aculeatus*) and coelacanth (*L. chalumnae*). **B,** Synteny plot of the short-beaked echidna (*T. aculeatus*) and platypus (*O. anatinus*) when compared to humans (*H. sapiens* / placental mammals), gracile agile opossums (*G. agilis* / marsupials) and chickens (*G. gallus* / reptiles). Class-level taxonomy (Sarcopterygii vs Actinopterygii) is displayed on the left, boxes with dotted lines indicates agastric species, coloured boxes indicate genes of interest, black boxes indicate pseudogene presence, numbers within a box indicates presence of only that family member and breaks or missing boxes in the syntenic block indicate gene absence.

We then explored the conservation and loss of genes implicated in gastric epithelial maintenance and development in jawed vertebrates. BLAST searching and synteny analysis of the sequences bearing gastrokine family homology (*Gkn1*, *Gkn2*, *Gkn3*), trefoil factor homology (*Tff1*, *Tff2*), as well as *Anxa10* and *Vsig1* indicated they were lost in agastric Actinopterygii taxa but conserved in the gastric grey bichir and spotted gar (Figure 2A).

**Figure 2:**
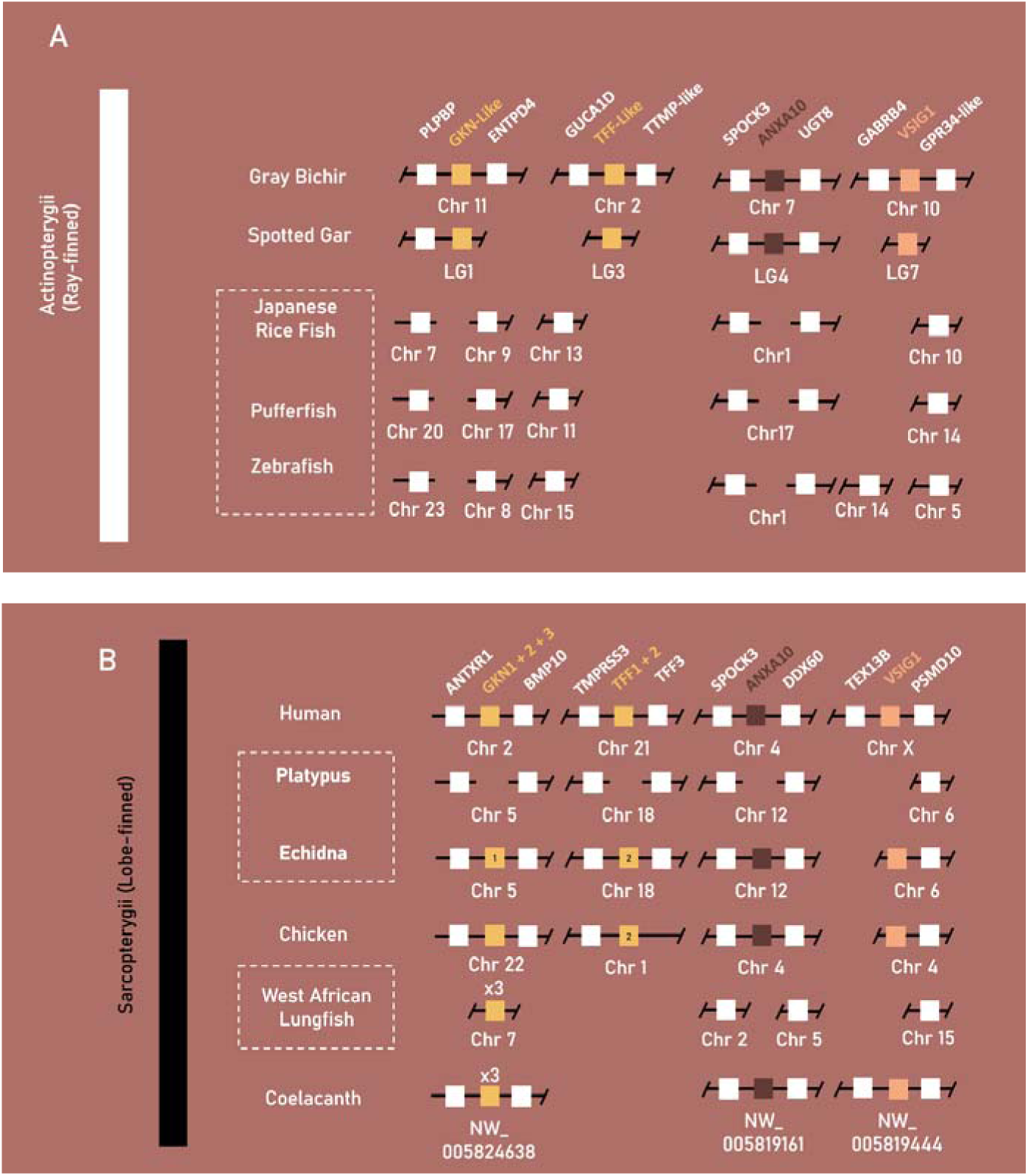
Synteny plots of genes implicated in gastric mucosal maintenance (*Gkn1*, *Gkn2*), gastric development (*Vsig1*) and unclear roles in gastric mucosal maintenance, growth and differentiation (*Anxa10*) in select vertebrate species. **A**, Synteny plot of Actinopterygii taxa: the agastric teleost species of the Japanese rice fish (*O. latipes*), pufferfish (*T. rubripes*) and zebrafish (*D. rerio*) compared to the gastric grey bichir (*P. senegalus*) and spotted gar (*L. oculatus*). **B** synteny plot of Sarcopterygii taxa: the agastric short-beaked echidna (*T. aculeatus*), platypus (*O. anatinus*) and West-African lungfish (*P. annectens*) compared to the gastric human (*H. sapien*), chicken (*G. gallus*) and coelacanth (*L chalumnae*). Class-level taxonomy (Sarcopterygii vs Actinopterygii) is displayed on the left, boxes with dotted lines indicates agastric species, coloured boxes indicate genes of interest, black boxes indicate pseudogene presence, numbers within a box indicates presence of only that family member and breaks or missing boxes in the syntenic block indicate gene absence.

In Sarcopterygii, gastrokine family paralogs (*Gkn1*, *Gkn2*, *Gkn3*) were mainly conserved together within a syntenic block flanked by *Antxr1* and *Bmp10*. Gastrokine paralogs 2 and 3 were lost in both monotreme species, but *Gkn1* was retained in the echidna. The three Gastrokine paralogs found in the West-African lungfish did not place within the conserved genomic region, and, further phylogenetic analysis of these sequences found that basal Sarcopterygii sequences (coelacanth, west-African lungfish) placed in a separate clade to all three vertebrate paralogs. Furthermore, the echidna *Gkn1* sequence displayed higher similarity to *Gkn3* sequences, and was placed within the vertebrate *Gkn3* clade with 100 percent bootstrap support (Figure S1).

Trefoil factor paralogs 1 and 2 (*Tff1*, *Tff2*) were absent in the coelacanth and West-African lungfish and found only in tetrapod species in a highly conserved syntenic block between *Tff3* and *Spock3* except in the platypus – missing both – and the echidna, which was missing *Tff1*. Genomic sequences of *Anxa10* and *Vsig1 w*ere found in all Sarcopterygii taxa but not in the platypus and west-African lungfish, confirming the association with the agastric phenotype found by Kato et al., (2024), (Figure 2B). These findings indicate that these genes evolved prior to the divergence of Actinopterygii and Sactoperygii, and were subsequently lost in agastric species.

### Sequence evolution and selection pressures in the gastrin release pathway between gastric and agastric jawed vertebrates (Gnathostomata)

We then investigated sequence evolution and Ka/Ks ratios in the gastrin release pathway in gastric and agastric vertebrates to see whether gastrin loss affected the evolution or selection of other pathway components.

The Actinopterygii clade of gastrin sequences displayed minimal difference in substitution per site between the gastric three-spined stickleback (0.48) and agastric Japanese rice fish (0.34),(Figure 3A). Branches leading to monotreme and therian mammal CCKBR clades displayed identical lengths (0.17 substitutions per site), but differed more post-divergence into the mouse and wombat (0.41 and 0.25 substitutions per site) when compared to the agastric platypus and echidna (0.07 and 2x10^-6^ substitutions per site). Actinopterygii CCKBRA sequences had similar substitutions per site within the clade of the pufferfish (0.15), the Japanese rice fish (0.15) and the three-spined stickleback (0.12). The pufferfish CCKBRB sequence contained a larger number of substitutions per site (0.2) when compared to three-spined stickleback (0.07) and Japanese rice fish (0.13), (Figure 3B).

**Figure 3:**
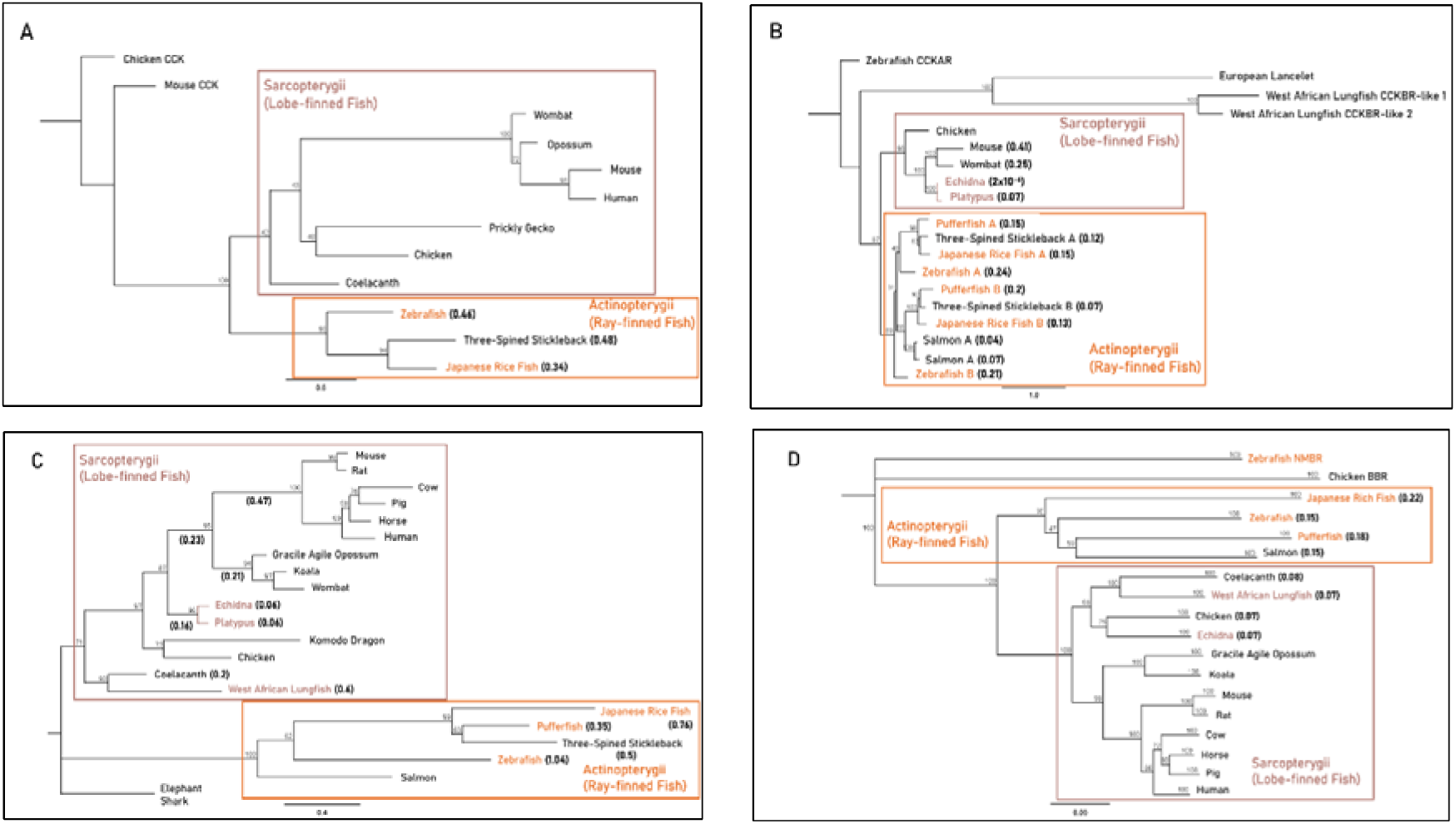
Maximum-likelihood amino acid phylogenies of the gastrin release pathway constituents: **A,** gastrin (GAST), **B,** cholecystokinin B receptor (CCKBR), **C,** gastrin-releasing peptide (GRP) and **D,** gastrin-releasing peptide receptor (GRPR) for select gastric and agastric vertebrate species constructed by the IQ-TREE web server with 1000 bootstrap replicates. Taxa, substitution models as chosen by Bayesian information criterion (BIC), outgroups and sequence references are in supplementary file 1. Paralogues of the CCKBR receptor for teleost lineages are identified with the letters A and B. Branch labels denote bootstrap percentage (%), coloured tip labels represent agastric lineages, coloured boxes outline class-level taxonomic groupings and branch length, scale bar and bracketed numbers measure substitutions per site to the nearest bifurcation.

In the Sarcopterygii clade, *Grp* sequences from the coelacanth and west-African lungfish displayed large variations in substitutions per site (0.2 vs 0.6). Though echidna and platypus sequences displayed identical estimated substitutions per site (0.06), there was substantial difference in branch length to the root of mammalian sequences between monotremes (0.16), marsupials (0.44) and placental mammals (0.7). In the Actinopterygii clade, branch lengths differed to the root of the clade with the Japanese rice fish (0.76), pufferfish (0.41) and three-spined stickleback (0.55), (Figure 3C). Sarcopterygii and Actinopterygii *Grpr* sequences little difference in substitutions per site between gastric and agastric species, as shown by comparisons between the salmon (0.15) and the pufferfish (0.18), and comparisons of the coelacanth (0.08) and west-African lungfish (0.07), (Figure 3D). These findings suggest acceleration of sequence evolution from gastrin release pathway constituents independent of gastric phenotypes (gastric versus agastric).

Finally, we calculated the ratio of synonymous to non-synonymous substitutions (Ka/Ks) using maximum-likelihood branch model selection analyses. We found overall that the *Gast, Cckbr*, *Grp* and *Grpr* genes were under purifying selection (Table S1). Foreground scenarios with significant likelihood-ratio tests (LRTs) in the *Cckbr* phylogeny included branches leading to the clade with Japanese rice fish, pufferfish and three-spined stickleback *Cckbrb* (*p* = 0.04, ω1 = 0.19, ω0 = 0.08), the three-spined stickleback *Cckbra* (*p* = 0.01, ω1 = 3x10^-3,^ ω0 = 0.07) and *Cckbrb* sequences (*p* = 1.12x10^-5^, ω1 = 0.33, ω0 = 0.08). In the *Grp* phylogeny significant LRTs included branches leading to the Tetrapod clade (*p* = 2.17x10^-5^, ω1 = 8.9x10^-4^, ω0 = 0.28), the mammalian clade (*p* = 0.03, ω1 = 0.02, ω0 = 0.24) and the mouse sequence (*p* = 0.04, ω1 = 0.02, ω0 = 0.24). The *Grpr* phylogeny included two significant LRTs; the branch leading to the pufferfish sequence (*p* = 0.02, ω1 = 0.11, ω0 = 0.04) and to the Sarcopterygii clade (*p* = 8.93x10^-7^, ω1 = 1.94x10^-3^, ω0 = 0.05), (Table 1). These findings indicate that independent bursts of sequence evolution in gastrin release pathway constituents consisted largely of conservative mutations with no direct correlation to gastric phenotype.

**Table 1:** Summary of significant findings from CodeML branch model selection analysis of *Cckbr*, *Grp* and *Grpr* sequences with columns for foreground branch (taxonomic grouping or species), the model-fitting metric (likelihood-ratio test *p*) and Ka/Ks ratio under differing foreground scenarios (foreground = ω1, background = ω0).

| Gene | Foreground branch | LRT<br>*p* value | Foreground<br>Ka/Ks ($\omega_1$) | Background<br>Ka/Ks ($\omega_0$) |
| --- | --- | --- | --- | --- |
| *Cckbr* | Japanese Rice Fish B, Pufferfish B and Three-Spined Stickleback B | 0.04 | 0.19 | 0.08 |
| *Cckbr* | Three-Spined Stickleback B | $1.12 \times 10^{-5}$ | 0.33 | 0.07 |
| *Cckbr* | Three-Spined Stickleback A | 0.01 | $3 \times 10^{-3}$ | 0.08 |
| *Grp* | Tetrapoda (Mammals and Reptiles) | $2.17 \times 10^{-5}$ | $8.9 \times 10^{-4}$ | 0.28 |
| *Grp* | Mammalia | 0.03 | 0.02 | 0.24 |
| *Grp* | Mouse | 0.04 | 0.06 | 0.24 |
| *Grpr* | Pufferfish | 0.02 | 0.11 | 0.04 |
| *Grpr* | Sarcopterygii (Mammals, Reptiles and Lobe-Finned Fish) | $8.93 \times 10^{-7}$ | $1.94 \times 10^{-3}$ | 0.05 |

## Discussion

Stomach anatomy in vertebrates can been broadly described as gastric or agastric. The latter phenotype lacks gross morphology, glandular epithelium and gastric acid secretion has been described in independent teleost lineages and monotremes. Changes in stomach morphology have been correlated with the loss of gastric enzymes (*Pga*, *Pgc*), proton pump subunits *(Atp4a*, *Atp4b*) and the gastrin hormone (*Gast*) in monotremes (Ordoñez et al., 2008; Castro et al., 2014; Zhou et al., 2021, Kato et al., 2024). Due to the complex constitution and function of the stomach – with a large complement of cell types, signalling pathways and physiological functions – we hypothesised that the gene losses associated with the agastric phenotype extended beyond these aforementioned losses. To test this, we investigated the evolution, selection and loss of the gastrin physiological pathway along with genes implicated in gastric development, and functionality in both gastric and agastric Sarcopterygii and Actinopterygii taxa.

### Gastrin physiological pathway: Correlations with gastric phenotype and genetic dispensability

We determined loss of gastrin in four of the five agastric species presented (platypus, echidna, pufferfish, West-African Lungfish) but surprisingly no substantial differences in substitution rate or selection pressures of the gastrin physiological pathway constituents (*Gast*, *Cckbr*, *Grp*, *Grpr*) between gastric and agastric species, or between species who had retained or lost gastrin. This conservation is likely due to diversification in functions and signalling pathways beyond gastric acid secretion (Li et al., 2003; Albalat & Cañestro, 2016).

The release of gastrin occurs via gastric distension or gastrin-releasing peptide receptor activation and potentiates the release of gastric acid (Ordoñez et al., 2008). Despite retaining the receptor, the zebrafish (*D. rerio*) has lost the genes necessary to release gastric acid (*Atp4a*, *Atp4b*) and most gastric morphology (Castro et al, 2014). In contrast to other agastric fish species – zebrafish (*D. rerio*) has retained gastrin. Recent research has found novel roles of gastrin and its related isoforms (progastrin and the glycine-extended gastrins) in cell proliferation, apoptosis and extracellular remodelling in colonic epithelium in a suite of vertebrate models (Dimaline & Varro, 2014). Furthermore, mouse gastrin knockout models present with heavily deleterious gastric phenotypes: achlorydic gastric environments with high incidences of gastric metaplasia, bacterial overgrowth, and tumours (Friis-Hansen, 2007). A low substitution rate for zebrafish *Gast* or the native receptor *Cckbr*, indicates that gastrin may have further diversified in functionality (perhaps due to the *Cckbr* duplication) since the third whole-genome duplication event (3R) in teleosts, and maintenance of these functions explains the sequence conservation (Brunet et al., 2006; Ozernyuk & Schepetov, 2022). This also suggests that loss of gastrin in other agastric species may not only be due to loss of gastric epithelium and gastric acid secretion.

Further, we found no association between gastric phenotype (gastric vs agastric) or loss of gastrin with the absence of *Grpr* in the agastric platypus and gastric three-spined stickleback. The gastrin-releasing peptide pathway is implicated not only in gastrin release, but also in food intake regulation and growth in fish models, and in sexual arousal, thermoregulation and circadian rhythm regulation in rodent models (Volkoff et al., 2005; Duan, Rico & Merchant, 2022; Oti & Sakamoto, 2023; Zhao et al., 2023). The loss of GRP-GRPR signaling despite a retention of gastrin the three-spined stickleback suggests either that gastric mechanical stimuli may be sufficient to potentiate gastrin release in this species or that an independent gastrin-release mechanism is yet to be explored. This independent mechanism may be partially explained by the fact that the GRP and GRPR peptides are from the “bombesin-like” gene family, a family of three homologous lineages which diverged before the advent of jawed vertebrates and all with various roles as both neuropeptides and gastrointestinal hormones. Though affinity is weak, there is agonism between GRP and the homologous receptors.(Baldwin, Patel & Shulkes, 2007; Jensen et al., 2008).

### Evolution of gastric genes implicated in the secondary loss of gastric phenotype in jawed vertebrates (Gnathostomata)

We noted the absence of gastrokine (*Gkn1*, *Gkn2*) and trefoil factor homologs (*Tff1*, *Tff2*), *Vsig1* and *Anxa10* in all agastric taxa to the exception of gastrokine-like sequences in the West-African lungfish. Convergent loss between agastric Actinopterygii taxa, the West-African lungfish and the platypus suggests a role in the secondary loss of stomach features in line with loss of gastric enzymes, proton pump subunits and gastrin (Ordoñez et al., 2008, Castro et al., 2014; Zhou et al., 2021; Kato et al., 2024). However, the echidna has an agastric phenotype and has retained *Gkn1*, *Tff2*, *Vsig1* and *Anxa10*. These findings align with our recent works in which the common monotreme ancestor likely lost antral glandular epithelium and pyloric restriction through *Nkx3.2* pseudogenisation and subsequent shift of developmental processes, but the echidna lineage displayed retention or re-establishment of pyloric-like restriction through potential compensation or an additional evolutionary event (Dann et al., 2024).

The presence of trefoil factor and gastrokine homologs in Actinopterygii taxa suggests the evolution of these genes around the divergence of jawed vertebrates (Gnathostomata), (Jiang & Applegate, 2011; Geahlen et al., 2013). Furthermore, presence of a homologous trefoil factor-like sequence in the vase tunicate (*Ciona intestinalis*) provides evidence of gene evolution in line with the advent of the alimentary canal in early ascidians (Table S1), (Nakayama & Ogasawara, 2017). Recent research has implicated these genes in the differentiation of squamous and glandular epithelium in the mammalian hindstomach as well as mucosal proliferation/differentiation and protection to chemical insult. The gastrokines and trefoil factor homologs, in particular, are known for their potent anti-tumorigenic properties and are implicated in several gastrointestinal cancers (Yan et al., 2011; Jahan et al., 2020).

Further *in-vivo* experimentation in a variety of biological models could determine whether these are newly-acquired functions or whether these features date back to early jawed-vertebrates (Oidovsambuu et al., 2011, Gerke et al., 2005; Tsai et al., 2015).

## Conclusion

Here, we investigated the evolution, selection and conservation of the gastrin physiological pathway and other gene involved in gastric function in gastric and agastric jawed vertebrates in order to further understand the independent evolutionary trajectories of the agastric phenotype. Interestingly, we discovered that the absence of a gastric gene repertoire (*Gkn1*, *Gkn2*, *Tff1*, *Tff2*, *Vsig1*, *Anxa10*) – which evolved prior to jawed vertebrate divergence – correlates with the agastric phenotype to the exception of the echidna, which may have retained some aspects of gastric functionality (Figure 4). We found that the loss of *Gast* and *Grpr* does not correlate with the gastric or agastric phenotype. Furthermore, we did not observe a shift in sequence evolution or selection associated with the loss of these genes in vertebrates. These findings differentiate aspects of the agastric phenotype and reveal convergent gene loss and retention in agastric species separated by more than 500 million years of evolution.

**Figure 4:**
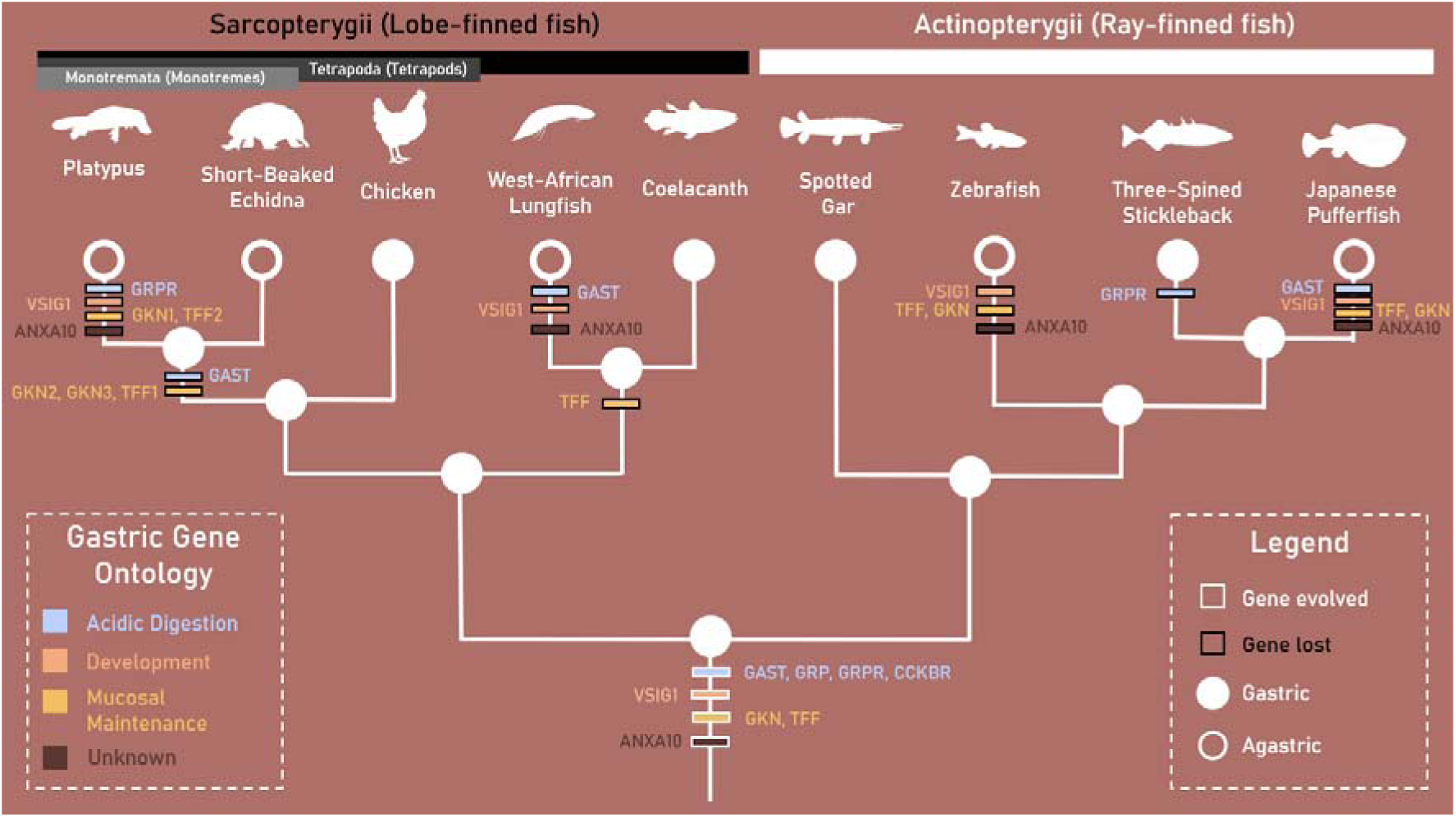
Phylogeny outlining the proposed gastric gene evolution, duplication and loss events between taxa displaying the gastric and agastric phenotype in Sarcopterygii (lobe-finned fish) and Actinopterygii (ray-finned fish) clades. Agastric taxa are denoted by hollow branch tips, gastric gene ontologies are outlined by gene colouration while gene evolution and loss events are signified by white and black borders respectively. Branches are not drawn to scale.

