## Supplementary Figure 1 for "Evolution of the Jawed Vertebrate (Gnathostomata) Stomach through Gene Repertoire Loss: Findings from Agastric Species"

Supplementary Materials:

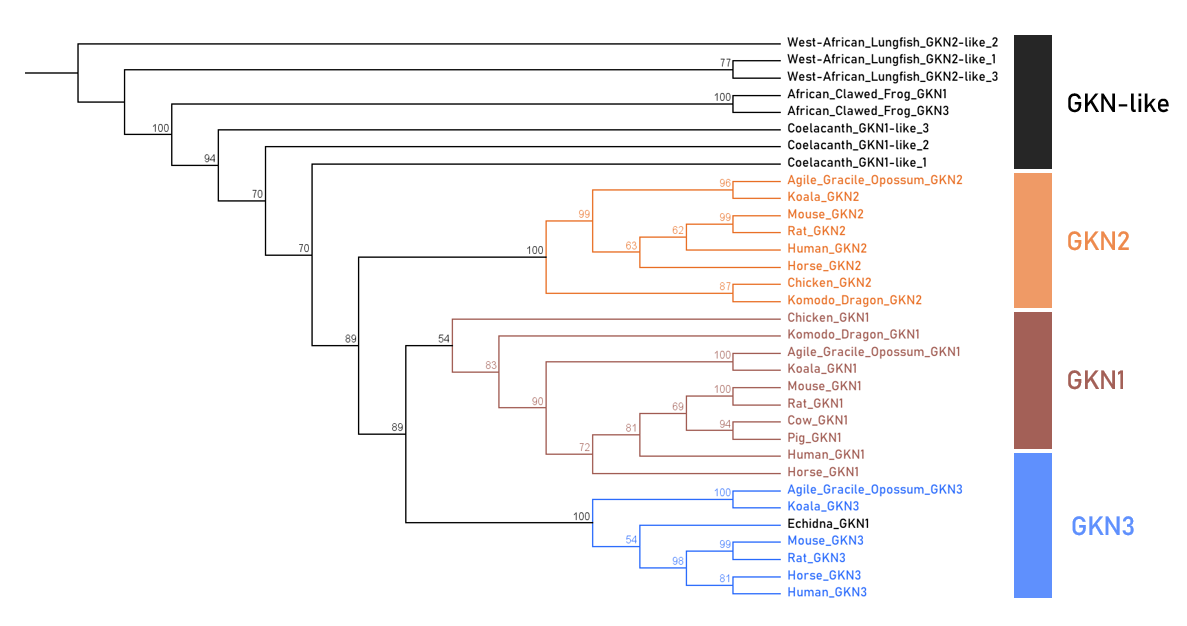

Figure S1: Maximum-likelihood cladogram of vertebrate gastrokine-like amino acid sequences constructed by the IQ-Tree web server using the JTT + G4 substitution model and 1000 bootstrap replicates. Branch values indicate bootstrap percentage and groupings of gastrokine-like sequences are superimposed on the right.
